# Preselection CD4^+^CD8^+^ thymocytes modulate TCR responsiveness following TCRβ selection

**DOI:** 10.1101/2025.10.03.680384

**Authors:** Esther Jeong Yoon Kim, Dominik A. Aylard, Zoë Steier, Daisy Ortiz, Jean Choi, Stephanie Chen, Aaron Streets, Nir Yosef, Ellen A. Robey

## Abstract

Modulation of TCR sensitivity during positive selection is critical to avoid negative selection and direct thymocytes into their appropriate lineage. Thymocytes just prior to positive selection (preselection) are highly responsive to low affinity self-ligands and are also actively rearranging their TCRα locus as they await a positive selection signal. Preselection DP thymocytes were thought to be relatively homogeneous and TCR modulation during this stage had not been previously described. Here we provide evidence for progressive gene expression changes within the preselection DP thymocyte population that correlates with a gradual loss of TCR responsiveness and a defect in upregulating TCR target genes associated with the CD4 fate. We relate these observations to the link between positive selection and T cell lineage commitment.

## Introduction

Signaling through the T cell antigen receptor (TCR) can lead to divergent outcomes including T cell differentiation, tolerance induction, survival, or death. The response depends not only on the nature of the peptide-MHC ligand and antigen presenting cell (APC), but also on the type of responding T cell (reviewed in ^1^). For thymocytes, successive stages of T cell development are accompanied by changes in TCR responsiveness to self-peptide-MHC ligands. For example, CD4^+^CD8^+^ (double positive or DP) thymocytes form multifocal, rather than condensed central immune synapses ^2,3^ and exhibit reduced calcium flux in response to TCR crosslinking compared to mature T cells ^4^. In addition, DP thymocytes that have not yet initiated positive selection (called preselection DP) are hypersensitive to low affinity ligands ^5,6^ a characteristic that allows them to initiate positive selection upon recognition of self-peptide MHC ligands on cortical thymic epithelial cells. In response to positive selection signals preselection DP downregulate their sensitivity to weak ligands and continue to dynamically regulate TCR responsiveness over the next few days as they undergo CD4 and CD8 lineage commitment (reviewed in ^7,8^). The mechanisms that modulate TCR responsiveness during thymic development and their impact on T cell fate remain poorly understood.

The somatic gene rearrangements that assemble the TCRα and β chains from gene segments are intertwined with T cell developmental progression and TCR signaling. At the early CD4^-^CD8^-^ (double negative or DN) stage, progenitor T cells first rearrange their TCRβ gene locus and production of functional TCRβ protein leads to signaling through the preTCR complex (TCRβ paired with the non-rearranging pTα chain) to drive proliferation and the transition from the DN to DP stage (reviewed in ^9^). After the TCRβ checkpoint, DP thymocytes stop proliferating and begin rearranging the TCRα locus. Thymocytes test their newly formed αβTCRs for the ability mediate positive selection, while also continuing to rearrange the TCRα locus ^10,11^. DP thymocytes have a lifespan of approximately three days, after which they either die or differentiate into mature CD4^+^CD8^-^ or CD4^-^CD8^+^ (single positive or SP) thymocytes ^12,13^ Because of successive TCRα gene rearrangements, “older” DP thymocytes harbor more distal rearrangements due to the replacement of initial V-J rearranged segments upon further rearrangement of upstream V and downstream J segments over time ^14–16^. It is generally assumed that any DP that rearranges an appropriate TCRα during the three-day time window is equally competent to undergo positive selection and give rise to CD4 or CD8 T cells.

Modulation of TCR responses during thymic development is also an important factor in CD4 versus CD8 T cell lineage commitment. During the first phase of positive selection, thymocytes undergo an initial wave of αβTCR signaling that activates general TCR target genes (e.g. *Egr1/2, CDd69, and Nr4a1/2*), as well as a program of CD4 gene expression including the transcription factors GATA3 and LEF1 that positively regulate expression of the CD4-defining transcription factor ThPOK, and CD5, a TCR tuning molecule that is preferentially expressed in CD4 compared to CD8 T cells ^17–21^. Thymocytes bearing TCRs specific for MHCII experience a relatively sustained 1^st^ TCR signaling wave and fully upregulate ThPOK, locking in the CD4 fate. In contrast, thymocytes whose TCR recognize MHCI experience a more transient 1^st^ TCR signaling wave, in part due to the loss of the CD8 co-receptor that accompanies the “CD4 audition”. This is followed by a second TCR signaling wave that serves as a quality control step during CD8 lineage specification. This second TCR signaling wave includes general TCR targets but lacks CD4 audition targets such as GATA3 and CD5 ^22^. It remains unclear why TCR targets associated with CD4 fate are activated in the early, but not the later TCR signaling wave during positive selection.

Here we focus on the preselection DP stage of development that follows TCRβ selection and precedes positive selection, providing evidence for phenotypic and functional heterogeneity within this population. We show that early preselection DP that have most recently completed TCRβ selection exhibit higher calcium flux upon TCR triggering and more robust induction of TCR target genes. We also show that resting calcium levels are relatively high in DN thymocytes, but gradually decrease throughout the preselection DP stage. We discuss the implications of these data for TCR repertoire formation and CD4 versus CD8 lineage commitment.

## Results

To investigate the thymocyte stages just prior to positive selection, we probed an existing single cell data set in which thymocyte populations from wild-type, MHC mutant, and TCR transgenic mice were analyzed by CITE-seq, providing transcriptomic and cell surface protein expression ^22^. While the previous study focused on populations undergoing positive selection via the αβTCR, the data set also included earlier DP populations, including DP that were proliferating following TCRβ selection, and two quiescent (non-proliferating) DP populations that did not show signs of αβTCR signaling (designated here as DP-Q1 and DP-Q2) (Fig. 1A). DP-Q1 and Q2 both display low expression of *Mki67* and *Cd5* genes (Fig. 1A), consistent with their lack of proliferation and TCR signaling respectively. Furthermore, DP-Q1 and Q2 both strongly express the *Rag1* gene as well as its positive regulator *Arpp21* ^22,23^ consistent with ongoing TCRα gene rearrangements prior to positive selection ^14,24,25^. Thus, DP-Q1 and DP-Q2 correspond to preselection DP thymocytes that have completed TCRβ selection but have not yet begun positive selection.

**Fig. 1.**
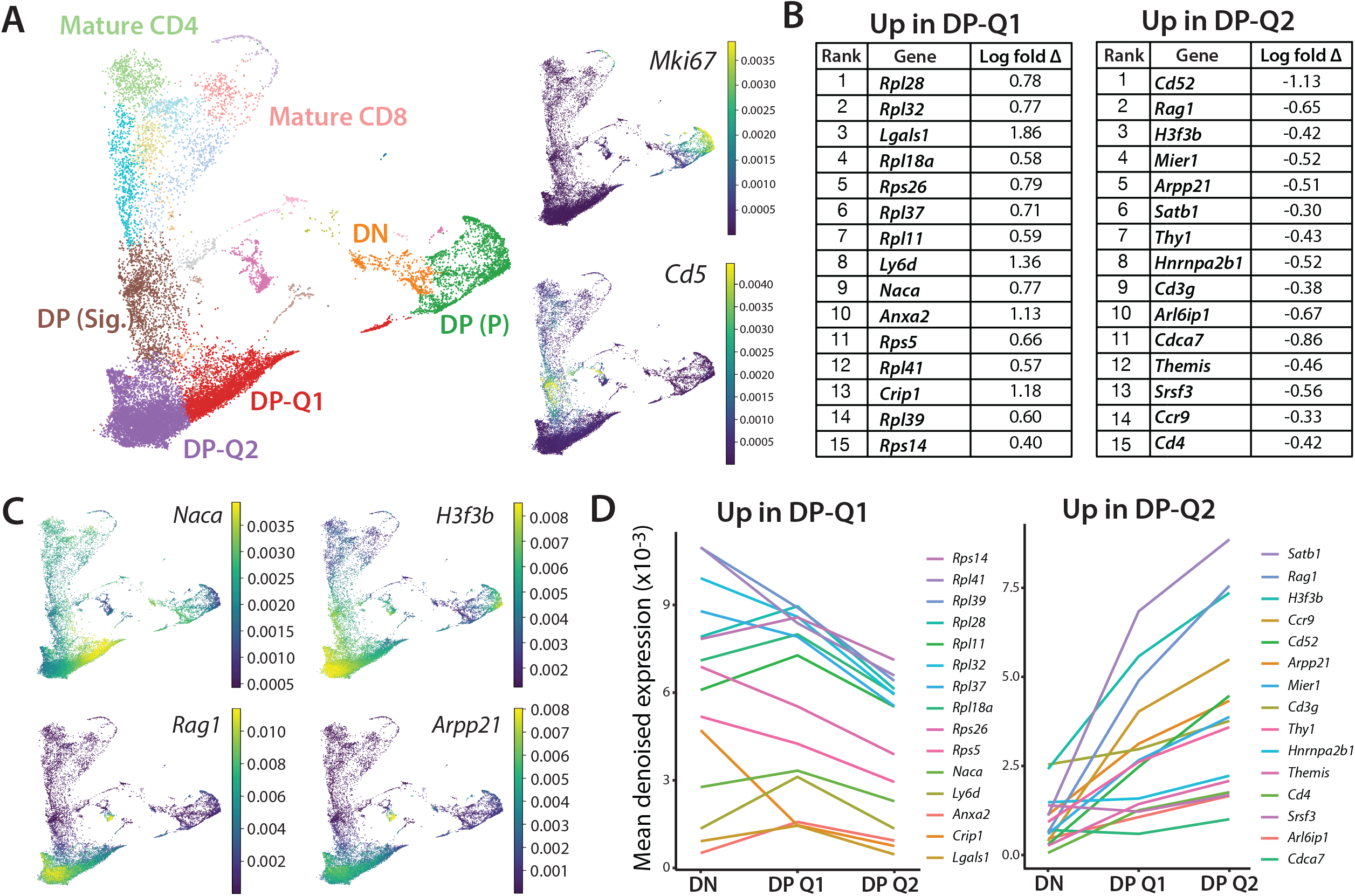
Differential gene expression between early and late preselection DP thymocytes in wild-type mice. **(A)** UMAP plot of totalVI latent space from CITE-seq dataset by Steier et al reveal distinct clusters of thymocytes during T cell development in wild-type B6 mice, labelled by cell type annotation ^22^. Four CD4^+^CD8^+^ double positive (DP) clusters are highlighted: DP(P) cluster represents proliferating DP thymocytes, DP-Q1 and Q2 represent early and late preselection DP thymocytes respectively, and DP (Sig.) represent signaled DP thymocytes. Right hand panels show denoised RNA expression of *Mki67* and *Cd5* across cell types. **(B)** Differential gene expression between DP-Q1 and DP-Q2 based on Wilcoxon rank-sum test. 15 genes with the most significant differential expression for each group are listed by rank. **(C)** Denoised RNA expression of selected genes that are differentially expressed between DP-Q1 and DP-Q2. **(D)** Change in mean denoised expression of genes listed in (B) compared between DN, preselection DP-Q1 and Q2 thymocytes.

We next used the CITEseq data set to explore transcriptional differences between DP-Q1 and DP-Q2 (Fig. 1B, C, Supplemental Fig. 1). Wild-type and TCR transgenic mice all contain DP-Q1 and DP-Q2 populations, although DP-Q2 is consistently less frequent in TCR transgenic mice, likely due to their preexisting TCR rearrangements that accelerate positive selection (Supplemental Fig. 1B). For simplicity, we focused on wild-type samples, and performed a Wilcoxon comparison between DP-Q1 and DP-Q2 (Fig. 1B). The 15 most significantly upregulated genes in DP-Q1, include ribosomal genes and the ribosome chaperone *Naca* ^26,27^ consistent with a more recent transition from proliferation to quiescence. On the other hand, DP-Q2 express higher levels of *Rag1* and *Arpp21*, consistent with their having spent more time in the quiescent preselection DP stage. DP-Q2 also have higher expression of chromatin regulators, including *H3f3b*, which encodes a histone variant involved in chromatin regulation ^15,28^, *Satb1* which encodes a chromatin organizer with key roles in thymocyte gene regulation ^29^ (Fig. 1C, Supplemental Fig. 1A). Many of the most significantly upregulated genes in DP-Q2 are lower in DN thymocytes, while there is a downward expression trend from DN to DP-Q1 for genes that are higher in DP-Q1 compared to DP-Q2 (Fig. 1D). Overall, the gene expression patterns of these populations suggest that both populations correspond to resting preselection DP undergoing TCRα rearrangements prior to positive selection, with Q1 and Q2 corresponding to earlier and later preselection DP, respectively.

To identify cell surface markers that distinguish between DP-Q1 and DP-Q2, and to further explore their identity, we compared protein expression between these populations from CITE-seq data (Fig. 2A, B). Notably, several proteins that are strongly expressed at the proliferating DP stage and downregulated upon transition to quiescence (e.g. CD304 (Neuropilin 1 or NRP1), CD71 (transferrin receptor 1), CD102 and CD54 are higher in Q1 compared to Q2. This is consistent with the interpretation that DP-Q1 transitioned more recently from the DN stage, compared to DP-Q2. We also identified CD150 (SLAM) and CD11a (integrin αL) as proteins that are more highly expressed on the surface of DP-Q2 cells.

**Fig. 2.**
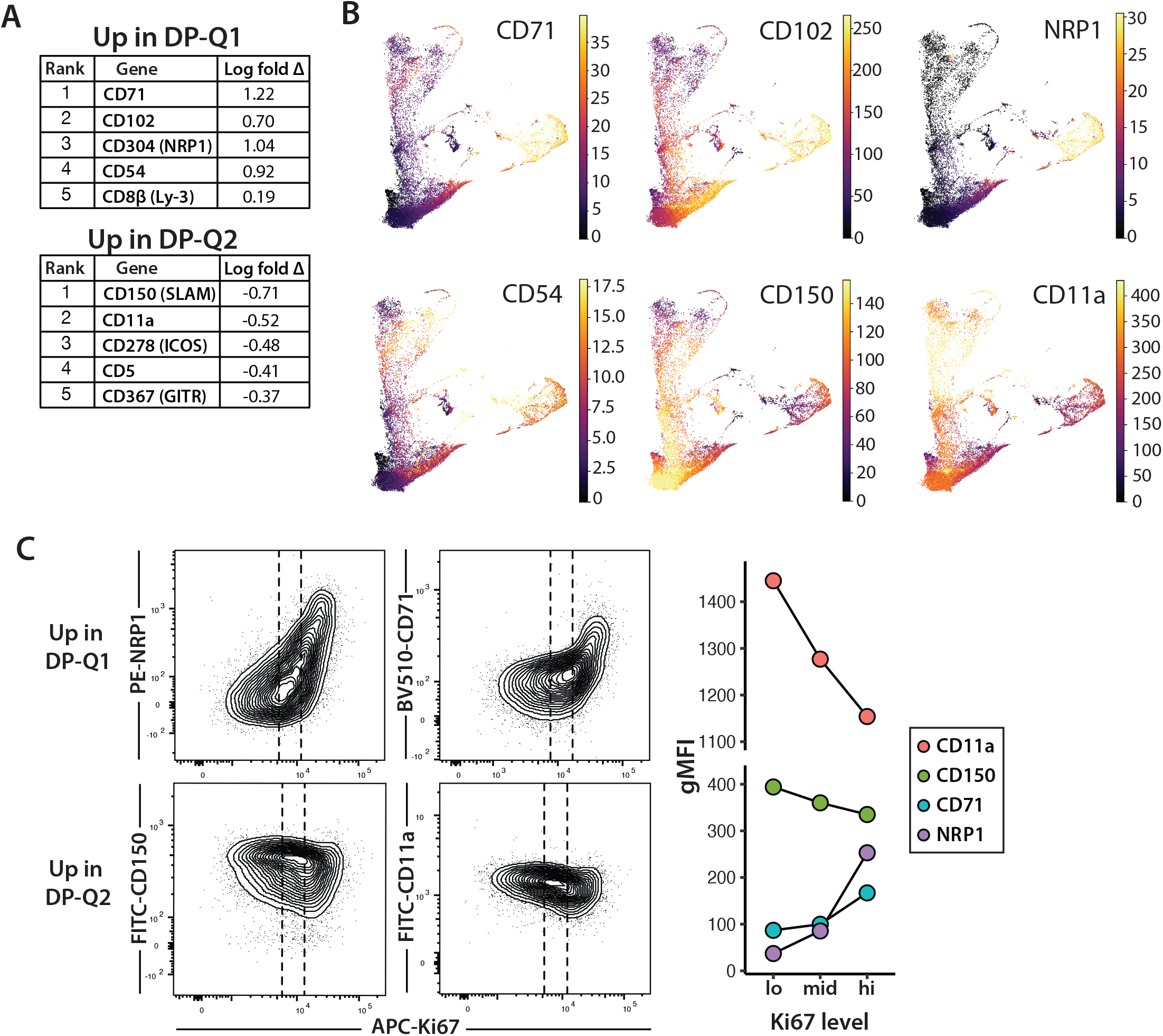
Cell surface markers distinguish early and late preselection DP thymocytes during thymic development. **(A)** Differential protein expression of between DP-Q1 and DP-Q2 amongst wild-type B6 thymocytes from Steier et al ^22^, based on Wilcoxon rank-sum test. Five proteins with the most significant differential expression are listed. **(B)** Denoised expression of proteins that distinguish DP-Q1 and DP-Q2 preselection thymocytes. Color intensity reflects protein expression across distinct thymocyte clusters. **(C)** Representative flow cytometry plots of Ki67 expression against surface levels of Q1/Q2 markers in preselection (CD5^lo^ CD69^lo^) DP thymocytes from wild-type B6 mice. To compare expression of Q1/Q2 markers within the non-proliferating preselection DP population, cells were divided into thirds according to Ki67 level (dotted lines; high, medium, low). gMFI within different Ki67 gates are plotted on the right. Data are representative of three independent experiments.

To confirm the relative age of DP-Q1 and Q2 populations, we performed flow cytometry combining cell surface markers with intracellular staining for the proliferation-associated protein, Ki67. Previous studies have shown that Ki67 protein decays slowly after cells exit cell cycle ^30^. Thus, Ki67 protein levels provide a measure of elapsed time since the last cell division. As expected based on previous studies ^31^, Ki67 expression levels were high in DN, proliferating and immature CD8SP (ISP), and proliferating DP thymocytes, low in mature SP thymocytes, and intermediate in DP, with signaled DP slightly higher than preselection DPs (Supplemental Fig. 2A). We then gated on preselection (CD5^lo^CD69^lo^) DP and compared Ki67 to surface markers identified using CITE-seq (Fig. 2C). As expected, CD11a and CD150 expression correlated inversely with Ki67, while NRP1 and CD71 show a positive correlation with Ki67 within the preselection DP population. We also observed that neonatal (day 3) mice have higher proportion (∼40%) of early preselection DP (Ki67^hi^, NRP1^hi^) thymocytes compared to ∼20% in adult mice (Supplemental Fig. 2B), consistent with the thymic accumulation of later preselection DP populations as mice age.

To confirm that the gene expression differences between earlier versus later preselection DP preceded, and were independent of, positive selection, we isolated preselection DP thymocytes from TCR transgenic mice crossed to a non-selecting MHC haplotype (AND TCR, *Rag1 KO*, H2^d^ and TG6 TCR, *Rag1 KO*, H2^b^, hereafter called pAND and pTG6). In these mice DP thymocytes are arrested at the preselection DP stage due to the absence of positive selecting MHC ligands. We observed a similar relationship between Ki67 and NRP1, CD71, CD150, and CD11a using preselection thymocytes from pAND and pTG6 mice (Supplemental Fig. 2C), confirming that these gene expression differences were not the result of early positive selection signals.

Given the differences in gene expression and age in the quiescent DP populations, we considered whether these differences might impact their ability to respond to TCR triggering. To investigate this question, we performed in vitro stimulations of preselection DP thymocytes from pTG6 mice using plate-bound TCRβ antibody. After 4-8 hours of stimulation, we read out TCR target gene induction using flow cytometry (Fig. 3A-C, Supplemental Fig. 3A). As expected, all preselection DP responded to TCR signaling as indicated by downregulation of surface TCRβ. However, we noted that target gene induction was consistently higher in earlier (Ki67^hi^) preselection thymocytes compared to later (Ki67^lo^) cells. This could be observed by a difference in the percentage of marker positive cells in response to TCR triggering (Fig. 3B, C), as well by the change in the overall intensity of marker expression within the population (gMFI) (Supplemental Fig. 3B). Moreover, while some targets were weakly induced in later preselection DP (e.g. CD5 and GATA3), others were less impacted by age of preselection DP (e.g. EGR2 and ID3). These data suggest the possibility that different branches of the TCR signaling pathway may be selectively impacted by the age of preselection DP thymocytes.

**Fig. 3.**
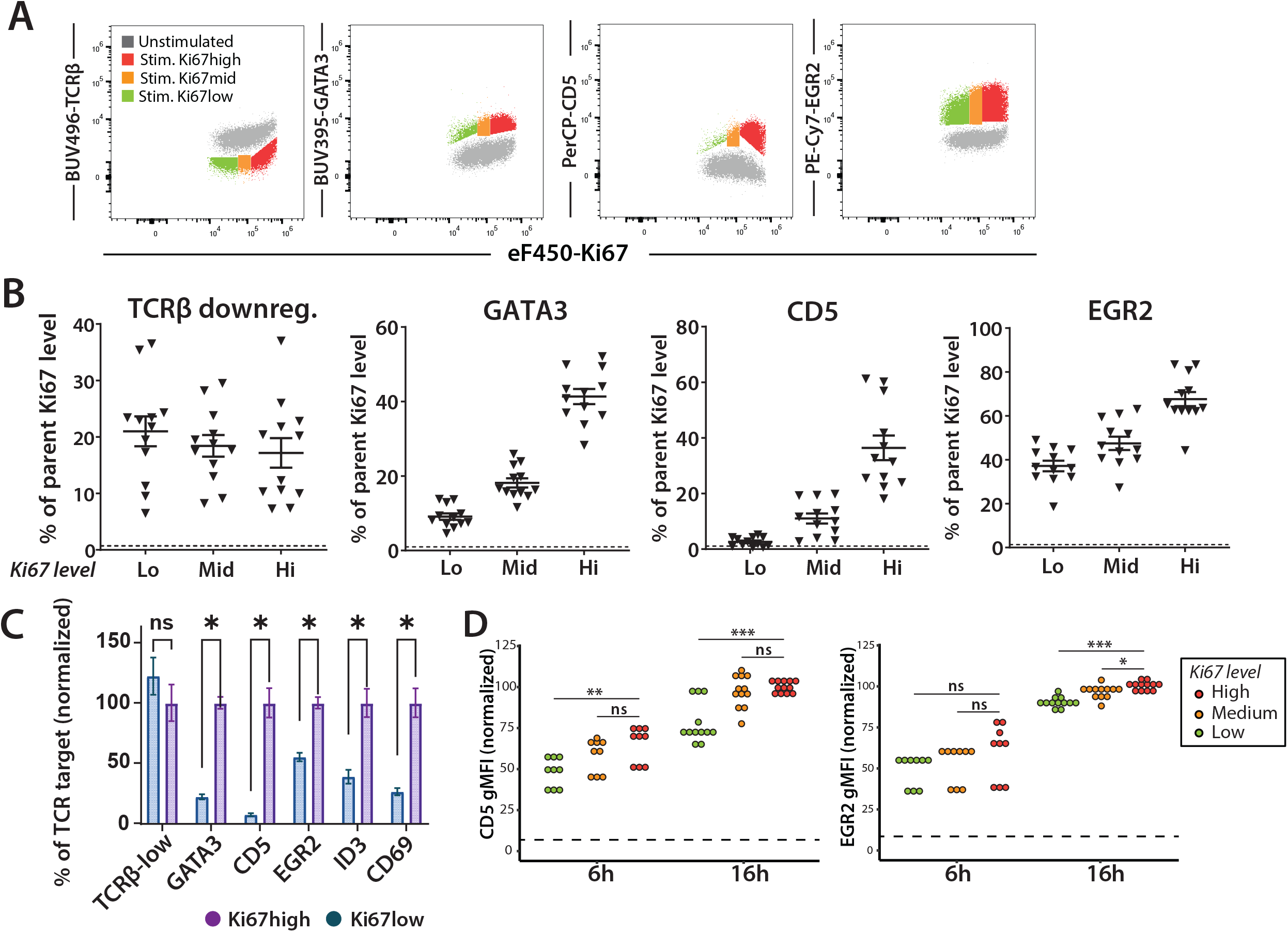
TCR target gene induction in early and late preselection DP thymocytes. **(A-C)** Preselection DP thymocytes from TG6 H-2^b^ *Rag2-/-* mice were stimulated via plate-bound αTCRβ antibody and marker expression was assessed by flow cytometry after 6 hours. (**A**) Representative flow cytometry plots of GATA3, CD5, EGR2 and TCRβ by Ki67 level. (**B**) Quantification of marker upregulation for different Ki67 levels. Background MFI was measured in the absence of stimulation, and these values are indicated by dashed line. Gating strategy is shown in Supplemental Fig. 3A. (**C**) Graph showing combined data for all markers normalized to the values of the Ki67^hi^ stimulated DPs. N= 4, performed in triplicates. Statistics via students t-test, between normalized percent TCR target value in Ki67^hi^ versus Ki67^lo^ DPs. * = p-value < 0.5. (**D**) Preselection DP thymocytes from MHC-deficient mice were stimulated via plate-bound α-TCRβ antibody and marker expression was assessed by flow cytometry after 6 and 16 hours. Gating strategy is shown in Supplemental Fig. 3C. gMFI values are normalized relative to Ki67^hi^ 16-hour stimulation condition for each marker. Background MFI was measured in the absence of stimulation, and these values are indicated by dashed line. N=3-4, p-values calculated by student’s t-test. *\* p<0*.*05, ** p<0*.*005, ***p<0*.*001*

To confirm these observations using non-TCR transgenic mice, we also performed in vitro stimulations of preselection thymocytes from MHC-deficient mice (Fig. 3D). Since some preselection DP in these mice do not yet express surface αβTCR, we gated on responding cells (marker positive) and used the intensity of marker expression (gMFI) within the positive gate as a measure of the response to TCR triggering (Supplemental Fig. 3C). Consistent with the data from pTG6 thymocytes, we observed more pronounced upregulation of CD5 and ERG2 in Ki67^hi^ (early) compared to Ki67^lo^ (late) cells. Moreover, this difference was more pronounced for CD5 compared to EGR2 (Fig. 3D). Thus, the increased responsiveness of early preselection DP is not a consequence of transgenic TCR expression but instead indicates that DP progressively alter their response to TCR stimulation over time after TCRβ selection.

The altered response to TCR triggering could impact how late vs early preselection DP respond to positive selection signals. To provide an in vitro readout of TCR responsiveness to relatively low affinity self-ligands that mediate positive selection, we examined responses to self-peptide MHC complexes displayed on bone marrow derived dendritic cells (BMDC) ^5,6,32,33^. For these experiments, we used preselection QFL thymocytes (QFL TCR+ β_2_M KO), cocultured them with BMDC from wild-type and MHC-deficient mice and measured upregulation of CD5, CD69, and EGR2 by flow cytometry after 6 or 16 hours (Fig. 4A, Supplementary Fig. 4).

**Fig. 4.**
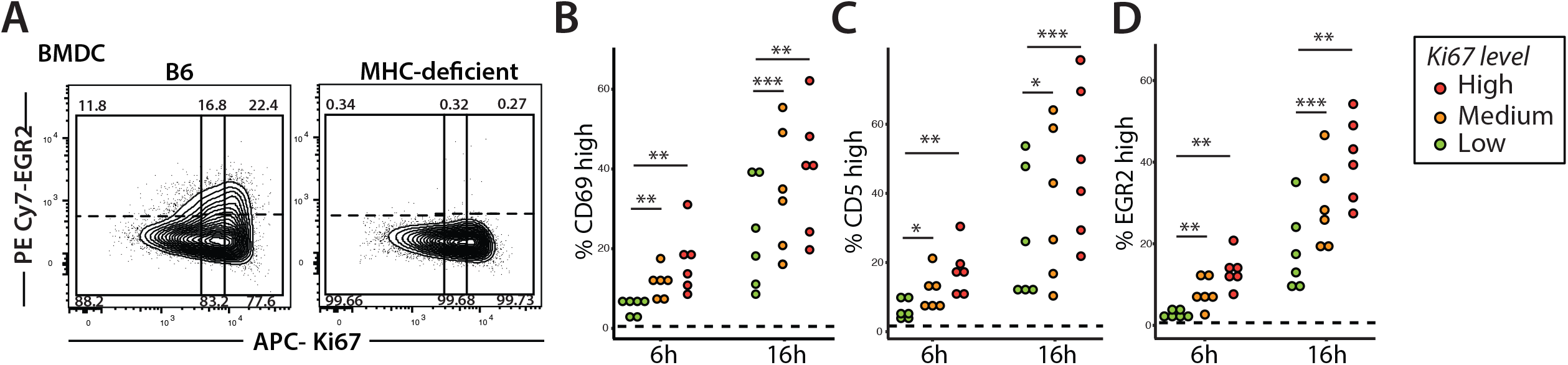
Early vs. late preselection DPs differ in their response to self-peptides. **(A)** Representative flow cytometry plots of EGR2 induction in preselection QFL thymocytes co-cultured with bone marrow derived dendritic cells (BMDCs) from either B6 or MHC-deficient mice (negative control for stimulation) at 16h. Preselection QFL thymocytes (Va3.2 + DP) are divided into thirds by Ki67 level (solid line; high, medium and low). Stimulation threshold is set by cells co-cultured with MHC-deficient BMDCs (dotted line), and assessed by upregulation of CD69 **(B)**, CD5 (**C)**, and EGR2 (**D)** within each Ki67 group. Each data point represents mean value from technical triplicates. N=6, p-values calculated by student’s t-test. *\* p<0*.*05, ** p<0*.*005, ***p<0*.*001*.

Consistent with the anti-TCR stimulation data (Fig. 3), earlier (Ki67^hi^) preselection QFL thymocytes exhibited stronger induction of downstream TCR signaling targets (Fig. 4B). Regardless of Ki67 levels, preselection QFL thymocytes did not respond to BMDCs derived from MHC-deficient mice, confirming that observed responses are MHC dependent.

To explore the mechanistic basis of the altered response of early vs late preselection DP to TCR triggering, we examined intracellular calcium flux, a central second messenger in the TCR signaling pathway (reviewed in ^34^). To this end, we used flow cytometry of thymocytes from pTG6 and pAND mice loaded with the ratiometric calcium indicator dye, Indo-1, together with CD3 crosslinking to mimic a strong TCR signal (Fig. 5, Supplemental Fig. 5). Since intracellular staining to detect Ki67 is incompatible with Indo-1 measurements, we used the cell surface markers NRP1 and CD150 to distinguish early vs late preselection DP thymocytes (Supplemental Fig. 5A). Calcium levels rapidly increased upon crosslinking, peaking within 3-4 minutes before gradual decline (Fig. 5A, B). For both pAND and pTG6 thymocytes, earlier preselection DP cells (NRP1^hi^) consistently exhibited higher levels of peak calcium compared to their later counterparts (NRP1^lo^) (Fig. 5C, D). Interestingly, prior to CD3 crosslinking, earlier DP (NRP1^hi^, CD150^lo^) thymocytes exhibit elevated baseline intracellular calcium levels compared to later DP (NRP1^lo^, CD150^hi^) (Fig. 5 E, F). Moreover, the populations just prior to preselection DP, DN and immature CD8SP (ISP) had even higher basal calcium, suggesting a gradual reduction in resting intracellular calcium correlating with thymocyte maturation (Fig. 5G and Supplemental Fig. 5B).

**Fig. 5.**
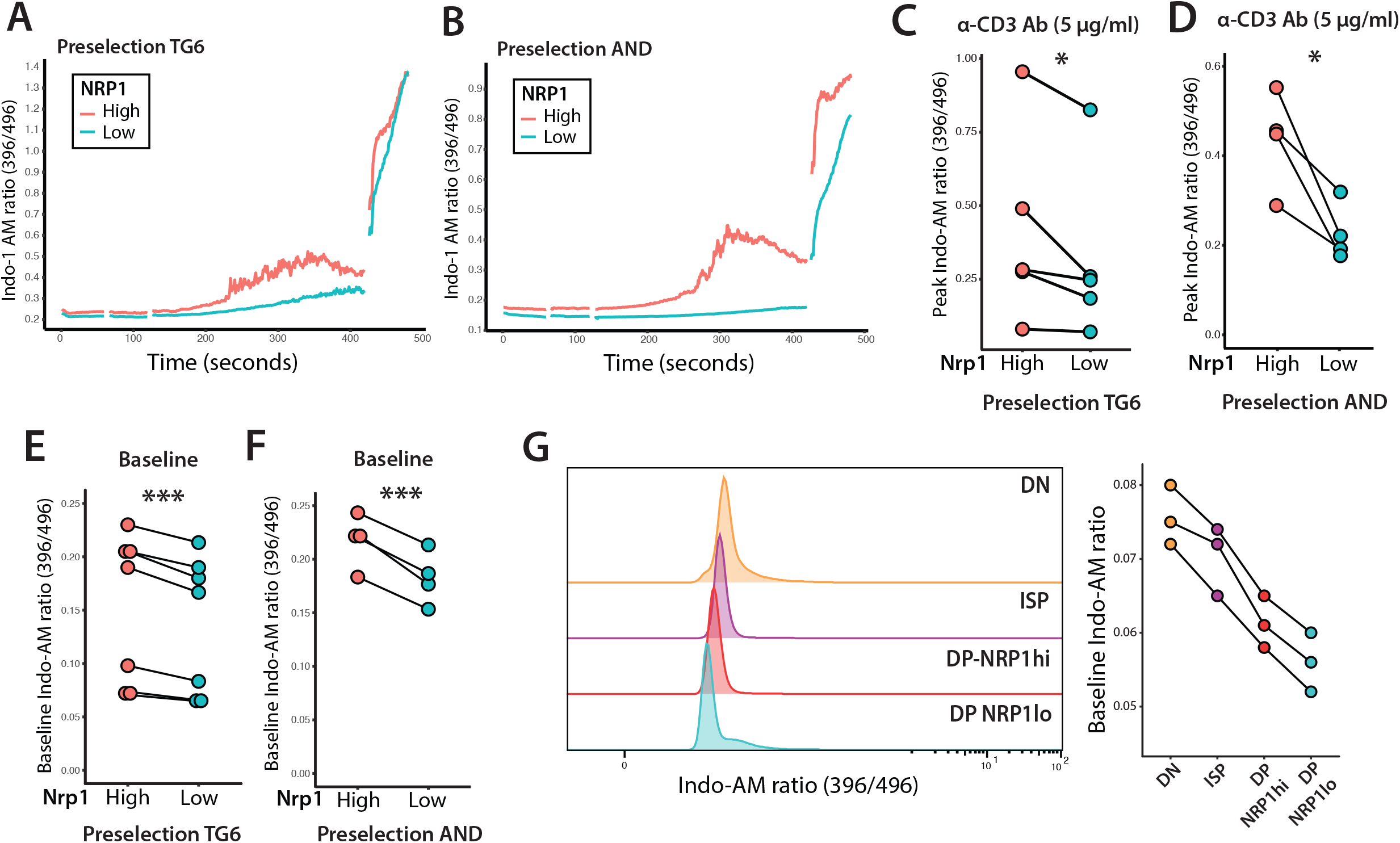
Intracellular calcium in early vs late preselection DP thymocytes. Preselection thymocytes from pTG6 (**A, C, E**) or pAND **(B, D, F)** mice were preloaded with Indo-1 and analyzed by flow cytometry. After collecting baseline values (1 minute), thymocytes were stimulated using anti-CD3ε antibody followed by anti-Armenian hamster IgG. Stimulation was assessed for 5 minutes, before addition of 1 μM ionomycin during the last minute of readout. **(A, B)** Representative plot of median Indo-1 ratio versus time in preselection thymocytes stimulated with 5ug/ml of anti-CD3ε antibody. **(C, D)** Comparison of peak Indo-1 calcium ratio between early (NRP1^hi^) and late (NRP1^lo^) preselection thymocytes in response to CD3ε-crosslinking. Each data point represents peak Indo-1 ratio value from an independent experiment. **(E, F)** Comparison of baseline Indo-1 calcium ratio in early (NRP1-hi) and late (NRP1-lo) preselection thymocytes. N=5 (pTG6), N=4 (pAND), p-values calculated by paired student’s t-test. *\* p<0*.*05, ** p<0*.*005, ***p<0*.*001*. **(G)** Representative histograms show baseline Indo-1 calcium ratio across DN, immature CD8SP (ISP), preselection NRP1^hi^ DP, and preselection NRP^lo^ DP thymocytes isolated from MHC-deficient mice in the absence of stimulation. Differences are quantified on the right (N=3).

## Discussion

Preselection DP thymocytes were previously thought to be a relatively homogeneous population with their ability to undergo positive selection primarily dependent on the expression of an αβTCR that confers the appropriate low to moderate reactivity for positive selecting ligands. Here we report unexpected phenotypic and functional heterogeneity within preselection DP, with earlier thymocytes (those that have most recently completed TCRβ selection) more poised to flux calcium and upregulate TCR target genes, and later DP thymocytes exhibiting higher *Rag* expression and a more limited ability to respond to TCR triggering. These data are relevant for understanding T cell fate decisions and TCRα repertoire selection, as we discuss below.

We recently presented a time-line of global gene expression changes during thymic positive selection ^22^ and used this information to formulate a unifying “Sequential Selection” model for CD4 versus CD8 lineage commitment ^7,8^ that incorporated elements of earlier Instructive, Stochastic/Selection ^35,36^, and Kinetic Signaling models ^37^. According to the Sequential Selection model, the initial stage of positive selection corresponds to a “CD4 audition” for both CD4-fated (MHCII specific) and CD8-fated (MHCI specific) thymocytes. The CD4 audition serves as the initial selection for a match between co-receptor expression and TCR specificity, since CD4-fated thymocytes experience stronger and more sustained signals due to dual recognition of MHCII by CD4 and TCR, whereas MHCI specific thymocytes fail the audition as they downregulate CD8. After a failed CD4 audition, MHCI specific thymocytes experience a second wave of TCR signal that overlaps with CD8 lineage specification. This second selection phase serves as a quality control step to ensure that only thymocytes bearing MHCI specific TCRs complete their maturation as CD8 T cells, while any MHCII specific thymocytes that failed to complete the CD4 audition are eliminated due to downregulation of CD4 and loss of TCR signal at this stage. Thus, the sequential selection model incorporates the notion, also invoked by the earlier stochastic/selection and kinetic signaling models, that the expression patterns of CD4 and CD8 coreceptors provides a selection against “co-receptor mismatched” thymocytes by the requirement for a TCR signal after co-receptor downregulation. This is consistent with experimental evidence for a role of co-receptor expression dynamics in lineage commitment, demonstrated most dramatically by genetically engineered “flipflop” mice, in which reversing CD4/CD8 expression patterns leads to CD4/CD8 lineage reversal ^8,38^.

Why do developing thymocytes first audition for the CD4 lineage before trying out the CD8 fate? We have previously proposed that this may relate to the higher average tonic TCR signals from self-recognition experienced by CD4 T cells compared to CD8 ^39^, perhaps due to different utilization of the Src family kinase LCK by CD4 versus CD8 ^40^. The current data showing that thymocytes that have most recently completed TCRβ selection are primed to receive relatively strong positive selection signals and to express genes associated with the CD4 fate (e.g. GATA and CD5), provides an additional explanation for why the CD4 fate is limited to the early phase of positive selection. Specifically, a thymocyte that expresses a TCR specific for MHCII soon after TCRβ selection may have a better chance of completing the CD4 audition compared to a thymocyte that has to try out one or more non-selecting TCRs before finally generating a TCR that recognizes MHCII. According to these arguments, CD4 T cells on average should be derived from earlier preselection DP compared to CD8 T cells and harbor “earlier” TCRα gene rearrangements (Suppl Fig 6). In support of this notion, previous studies have reported that CD4 SP use more proximal TCRα to TCRJ segment rearrangements compared to CD8 SP in both mice ^41^ and humans ^42^.

What are the molecular mechanisms responsible for the gradual change in TCR response as preselection DP age? While there are significant changes in mRNA and protein expression between early versus late preselection DP thymocytes, these differences are subtle compared to the large-scale gene expression changes that accompany TCRβ selection and positive selection, raising the possibility that non-transcriptional changes may contribute to the altered TCR responses. In this regard, it is intriguing that resting cytosolic calcium levels are relatively high in DN thymocytes and gradually decrease with progressive stages of maturation, including as preselection DP rearrange their TCRα locus and await a positive selection signal. Moreover, TCRβ selection is accompanied by cell proliferation and high metabolic activity, and the transition to full quiescence may be accompanied by gradual changes in signaling molecules and metabolic intermediates as these decay after TCRβ selection. Thus, we speculate that the relatively higher levels of signaling molecules and metabolic intermediates may prime early preselection DP to undergo positive selection and give rise to CD4 T cells. On the other hand, while later preselection DP thymocytes retain the ability to upregulate many TCR target genes, their fate may be skewed toward becoming a conventional CD8 T cell or undergoing agonist selection to give rise to interepithelial lymphocyte precursors upon receipt of a TCR signal (reviewed in ^43^)

In summary, our study shows that preselection DP thymocytes exhibits phenotypic changes and reduction in TCR reactivity as a function of elapsed time since TCRβ selection. A better understanding of how these changes impact different branches of the TCR signaling pathway, and their impact on CD4/CD8 differential potential remain areas for future investigation.

## Materials & Methods

### CITE-seq data analysis

Thymocyte CITE-seq data and metadata were accessed via zenodo (https://zenodo.org/records/12583813) and supplementary data files ^22^. Cells were filtered to exclude doublets and exclude samples sorted by FACS for positive selection. Cells were further filtered for wild-type (B6) samples for all analyses except cell type abundance analyses, which included all unsorted wild-type and transgenic samples. Cell type abundances were calculated per mouse. For protein data, pseudo-bulk differential expression analysis was conducted using decoupler ^44^ with PyDESeq2 ^45^. Due to the low expression levels of RNA relative to protein, RNA differential expression testing was conducted by Wilcoxon rank-sum test using Scanpy ^46^.

### Mice

All mice were maintained and bred under pathogen-free conditions in accordance with guidelines approved by the Institutional Animal Care and Use Committees at the University of California, Berkeley. C57BL/6 (WT) mice were obtained from The Jackson Laboratory. Preselection TG6 *Rag1-/-* ^47^ and AND *Rag1-/-* ^48^ thymocytes were generated by crossing each TCR transgenic onto a non-selecting background, MHCI KbDb and MHCII I-Ad, respectively. Preselection QFL thymocytes were generated by crossing previously described QFL transgenic mice ^33^ onto MHCI-deficient background (*B2m*-/-obtained from Taconic). MHC-deficient animals were generated by intercrossing *B2m*-/- and I-Aβ-/- mice ^49^.

### Thymocyte isolation

Thymic lobes were dissected from euthanized mice and mechanically dissociated in homogenizers. Cells were strained through 70μm nylon mesh filters to obtain single cell suspension. Following centrifugation, RBCs were removed in ACK buffer, and thymocytes were washed before resuspension in complete RPMI (cRPMI: 10% FBS, 8mM HEPES, 1mM sodium pyruvate, 55 μM β-mercaptoethanol, non-essential amino acids, L-glutamine, penicillin-streptomycin) for subsequent use.

### Flow cytometry

Isolated thymocytes were stained using the following antibodies: CD4 (clone RM4-5 BD Biosciences), CD8a (clone 53-6.7, BD Biosciences), CD5 (clone 53-7.3, BD Biosciences), CD69 (clone H1.2F3, BioLegend), TCRβ (clone H57-597, BioLegend), NRP1 (clone 3E12, BioLegend), CD150 (clone mShad150, Invitrogen), CD11a (clone M17/4, eBioscience), CD71 (RI7217, BioLegend), TCR Vα3.2 (clone RR3-16, BioLegend),Ki67 (clone B56, BD Biosciences), EGR2 (clone erongr2, eBioscience), GATA3 (clone L50-823, BD Biosciences). Cells were stained in 2.4G2 supernatant with antibodies against surface proteins for 30 minutes at 4°C. Subsequently, cells were stained with PBS containing viability dye, either Zombie NIR (BioLegend 423105) or GhostDye (Cell Signaling Technologies 598635), for 15 minutes at 4°C. For staining for intracellular proteins, cells were fixed and permeabilized using the FoxP3/Transcription Factors Staining Buffer Set (Invitrogen 00-5523-00). Analysis was performed on BD Symphony A3 or BD LSR Fortessa X20 or Cytek Aurora.

### TCRβ antibody-mediated stimulation

48-well plates were coated with 150μL solution of TCRβ antibody (Invitrogen 14-5961-85) diluted in PBS to a concentration of 5μg/ml. For unstimulated controls, wells were coated with equal volume of PBS only. These were incubated in a 37°C cell culture incubator for 1 hour. Antibody solution and PBS were aspirated and replaced with 500μL cRPMI . Half a million preselection thymocytes were added per well and stimulated at 37°C. Cells were stained for TCR signaling markers and stimulation was assessed using flow cytometry.

### Bone marrow dendritic cell generation and in vitro stimulation

On day 0, harvested bone marrow cells were subject to red blood cell (RBC) lysis in ACK buffer (0.15M NH_4_Cl, 1mM KHCO3, 0.1mM Na_2_EDTA) for 5 minutes at room temperature. In each well of a 6-well plate, 3 x 10^6^ cells were cultured in complete DMEM containing 20ng/ml GM-CSF (Peprotech #315-03). Media changes were performed at days 2 and 4 with cDMEM freshly Supplemented with GM-CSF, before collection with 10 minute EDTA treatment at day 6. For in vitro stimulation, 250-300K BMDCs were plated in each well of a 48-well plate. BMDCs were given 30-60 min to settle to the bottom of the plate, and half a million preselection thymocytes were added. Cells were collected after 6 and 16 hours of co-culture and marker expression was assessed using flow cytometry.

### Indo-1 calcium assay

20-30 million freshly prepared thymocytes were resuspended in 1ml of cRPMI with 2μM Indo-1AM (Invitrogen I1223). After 30-minute incubation in at 37°C protected from light, cells were split between FACS tubes (8-10 million cells/tube), washed with cRPMI, and stained with appropriate surface antibodies followed by viability dye. Washed cells were resuspended in 200μl cRPMI and kept on ice until the flow cytometer was ready. Shortly before running samples on the cytometer, each tube of cells is briefly warmed up on a 37°C heat block. After 1 minute of acquisition of baseline Indo-1 ratio, 5 μg/ml of α-CD3ε antibody (Invitrogen 14-0031-86) is added to tube, briefly vortexed, and another minute of acquisition is followed by addition of 5 μg/ml AffiniPure goat anti-Armenian hamster IgG (Jackson Laboratories 1277-005-099). After 5 more minutes, 1 μM ionomycin and acquisition is continued for an additional minute.

## Acknowledgements

We thank Y. Nakao, M. Colden, and K. Ortega for technical assistance and provision of materials for experiments. We thank the Cancer Research Lab Flow Cytometry Core Facilities at UC Berkeley, including K. Heydari, M. Delcroix, and H.Dhaliwal for their help operating the flow cytometers.

