## Supplemental Figures for "Preselection CD4^+^CD8^+^ thymocytes modulate TCR responsiveness following TCRβ selection"

**Supplemental Fig. 1**

**(A)** Denoised RNA expression of genes that distinguish DP-Q1 and DP-Q2 preselection thymocytes. Color intensity reflects relative gene expression across distinct thymocyte clusters.

**(B)** DP-Q1 and DP-Q2 cell type abundance per mouse compared across wild-type (B6) and transgenic TCR animals. Each dot represents one mouse.

**Supplemental Fig. 2**

**(A)** Representative gating strategy to identify thymocytes at different developmental stages. DN thymocytes were further gated for TCRβ$^{lo}$ cells. DP thymocytes were divided into signaled ("DP-sig") and preselection populations based on expression of CD5 and CD69. Among preselection DP thymocytes, those with high forward scatter were further distinguished as proliferating DP ("DP-prolif"). Immature CD8 single positive (ISP) cells were distinguished from mature CD8SP cells based on low levels of surface TCRβ. For each developmental subset, Ki67 levels are shown as histograms.

**(B)** Representative flow cytometry plots of DP thymocytes harvested from wild-type B6 neonates (day 3 and day 10) and adult mice. NRP1 expression was examined within preselection (CD5$^{lo}$ CD69$^{lo}$) DP thymocytes, and % of NRP1$^{hi}$, Ki67$^{hi}$ cells as well as NRP1 gMFI were quantified. Each dot is a value from an individual mouse. 1-3 mice were sacrificed for each time point for every experiment, N=3, p-values calculated by student's t-test. * *$p<0.05$*, ** *$p<0.005$*, ****$p<0.001$*

**(C)** DP-Q1/Q2 marker expression across Ki67 levels in DP thymocytes harvested from TG6 and AND TCR transgenic mice on non-selecting backgrounds (AND TCR, H2$^{d}$, *Rag1-/-* (pAND) and TG6 H-2$^{b}$ *Rag2-/-* (pTG6).

**Supplemental Fig. 3**

**(A)** Gating strategy for pTG6 thymocytes to assess TCR target induction

**(B)** Graph showing combined data for all markers normalized to the gMFI of the Ki67high stimulated DPs. N=4 performed in triplicates, each dot is a technical replicate.

**(C)** for preselection MHC-deficient thymocytes to assess TCR target induction.

**Supplemental Fig. 4.**

Gating strategy for preselection QFL thymocytes to assess BMDC-mediated stimulation.

**Supplemental Fig. 5.**

**(A)** Gating strategy to assess identify early (NRP1$^{hi}$) and late (NRP1$^{lo}$ CD150$^{hi}$) preselection TG6 and AND thymocytes for calcium measurements.

**(B)** Gating strategy to separate DN, preselection NRP1$^{hi}$, and preselection NRP1$^{lo}$ thymocytes from MHC KO mouse to compare baseline calcium levels. Representative plot of Indo-1 ratio in the absence of stimulation shown.

**Supplemental Figure 6**

Working model: Prior to the DP stage, DN thymocytes (red circles) rearrange their TCRβ genes, and in frame rearrangements trigger preTCR signaling, proliferation, and the DN to DP transition (yellow circle). Preselection DP thymocytes (green circles) undergo successive rearrangements of the TCRα locus, while testing their receptors for the ability to recognize positive selecting ligands. At the early preselection stage (DP-Q1), TCR recognition leads to more robust positive selection signals, enhancing the efficiency of CD4 lineage commitment for thymocytes bearing

MHCII specific TCRs. Thymocytes that spend longer at the preselection DP stage experience weaker positive selection signals making them less able to complete the CD4 audition, while retaining the ability to later develop as CD8 T cells if they rearrange a TCR specific for MHCI. The resulting prediction, that CD4 T cells would harbor more proximal TCR$\alpha$ rearrangements than CD8 T cells, is supported by previous studies (Park et al. 2020b; Lu et al. 2023).

### Suppl fig 1

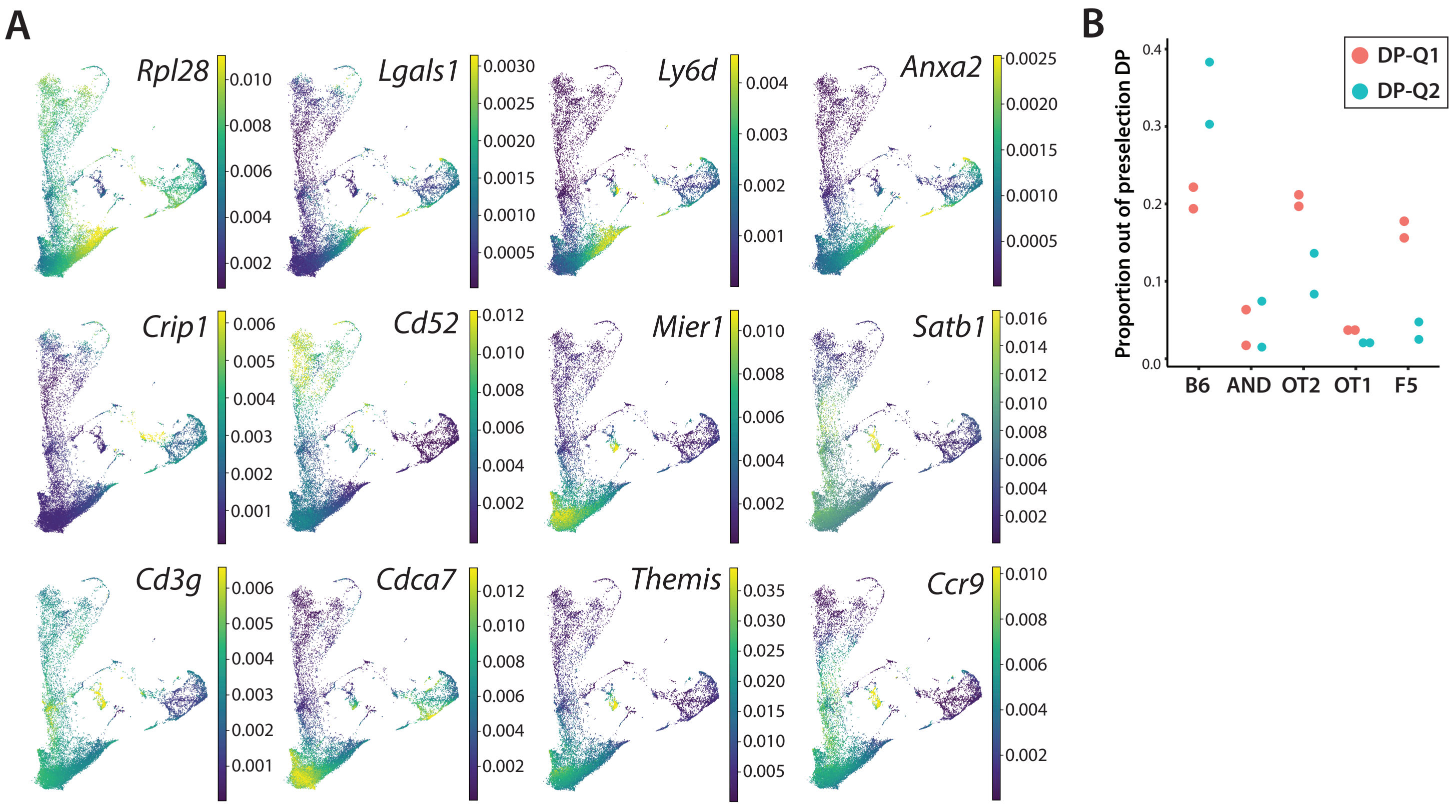

A
Rpl28
Lgals1
Ly6d
Anxa2
Crip1
Cd52
Mier1
Satb1
Cd3g
Cdca7
Themis
Ccr9
B
Proportion out of preselection DP
DP-Q1
DP-Q2
B6
AND
OT2
OT1
F5

#### Suppl. figure 2

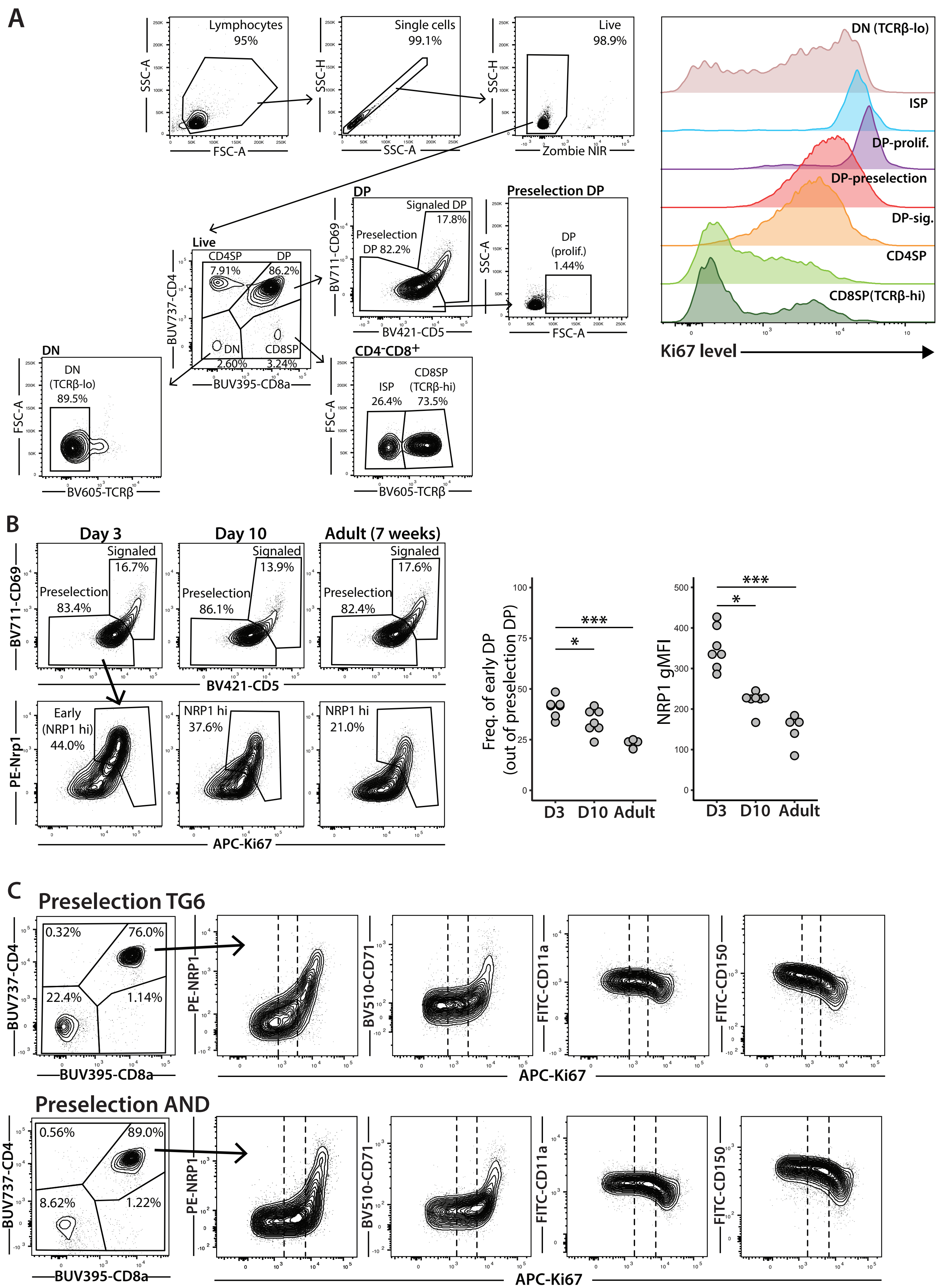

#### Suppl. figure 3

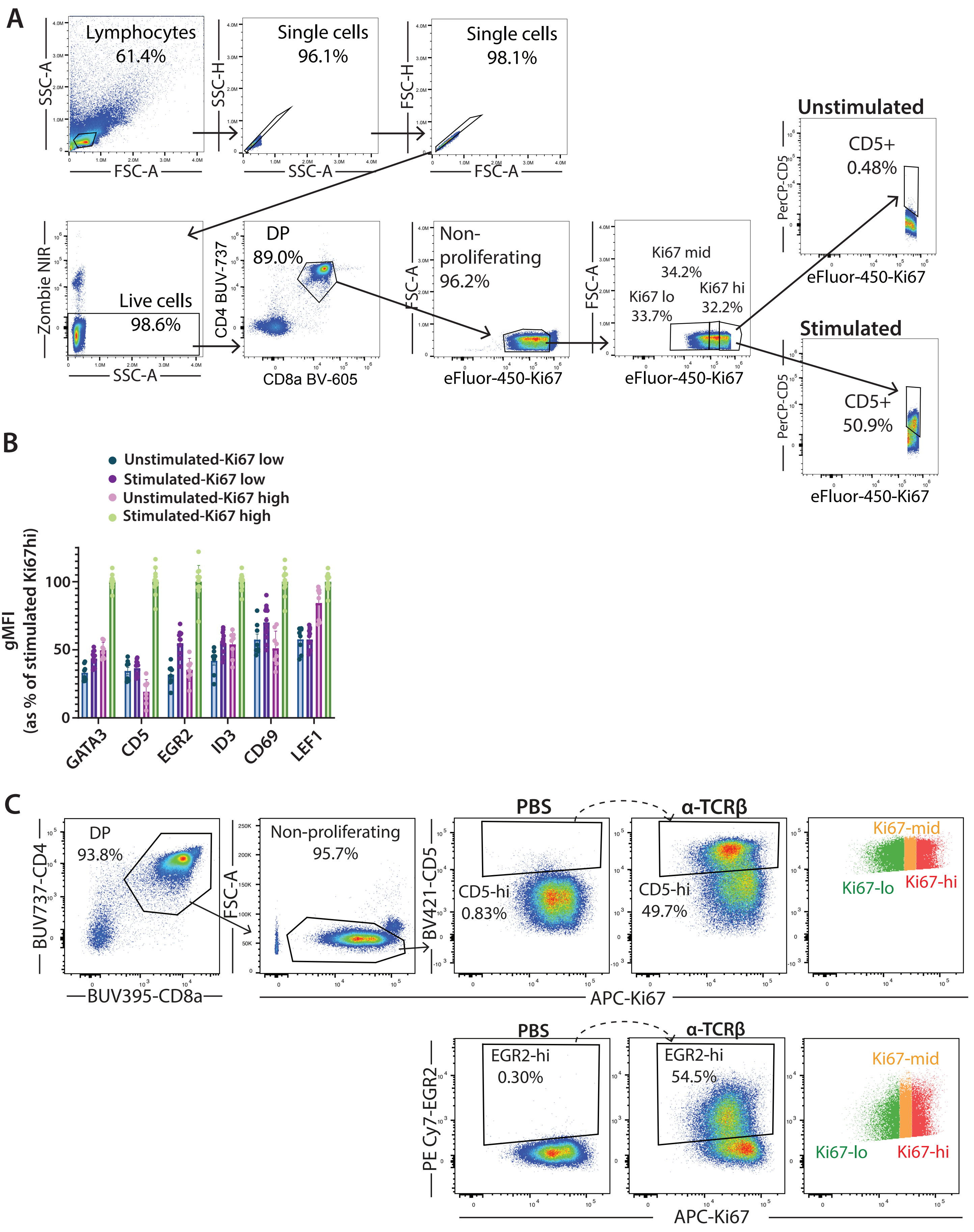

**Suppl. figure 4**

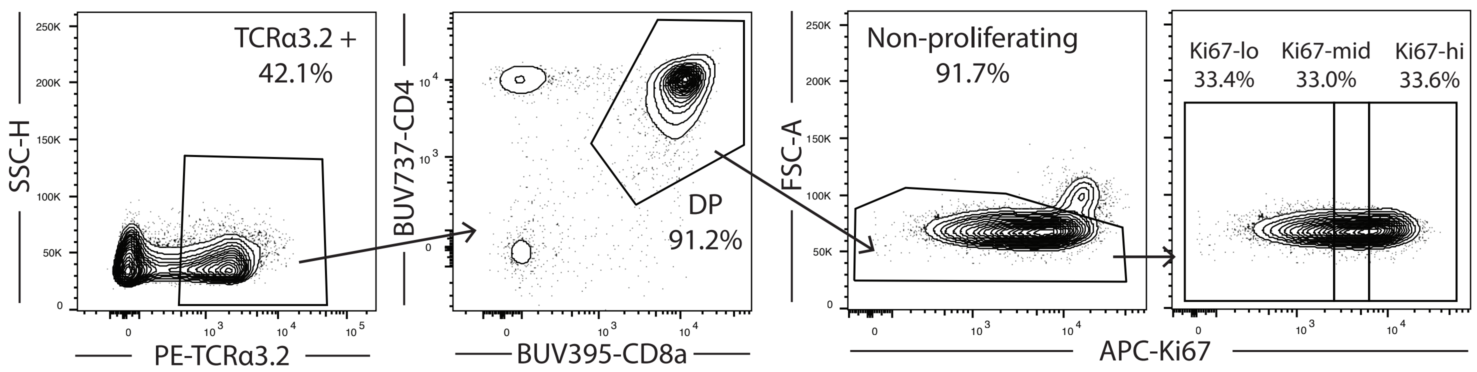

#### Suppl. figure 5

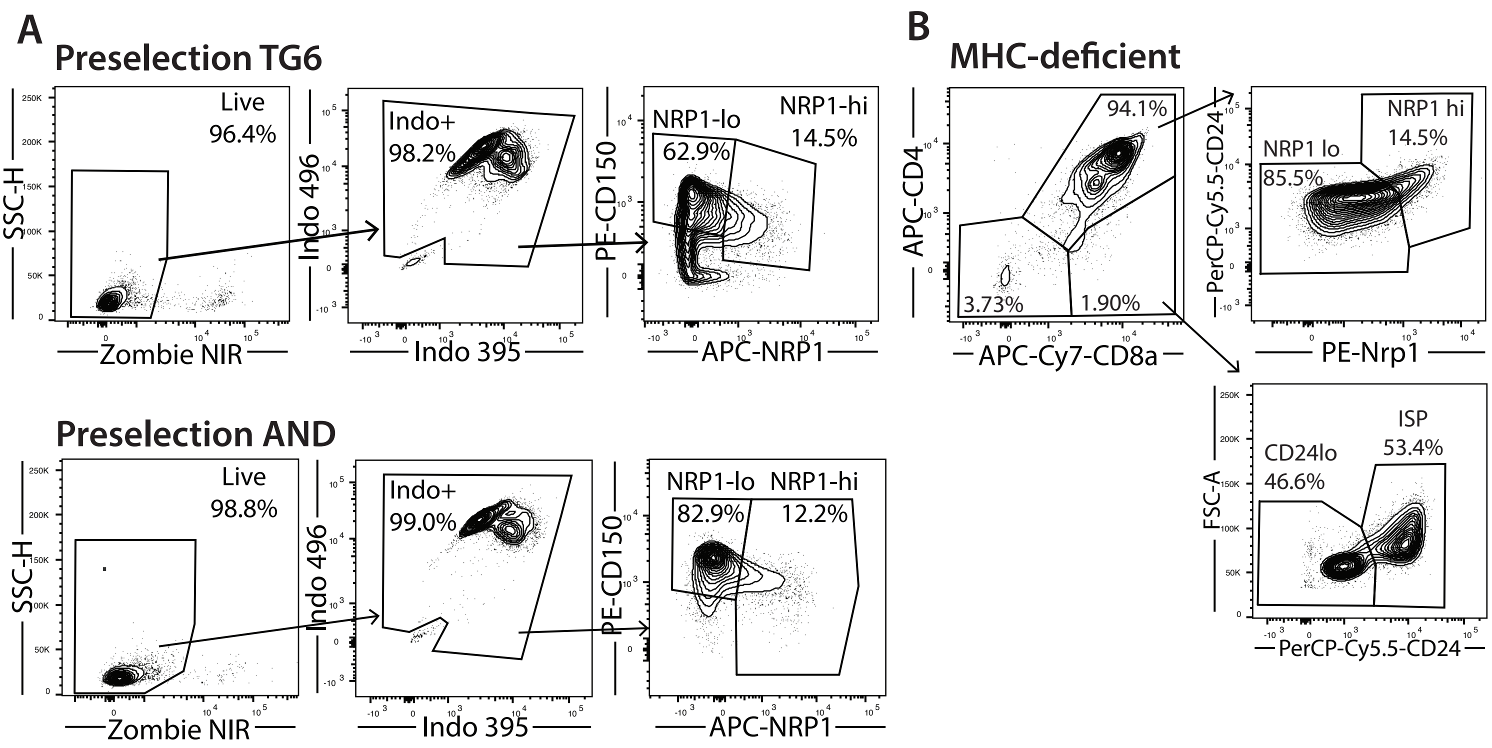

#### Suppl. figure 6

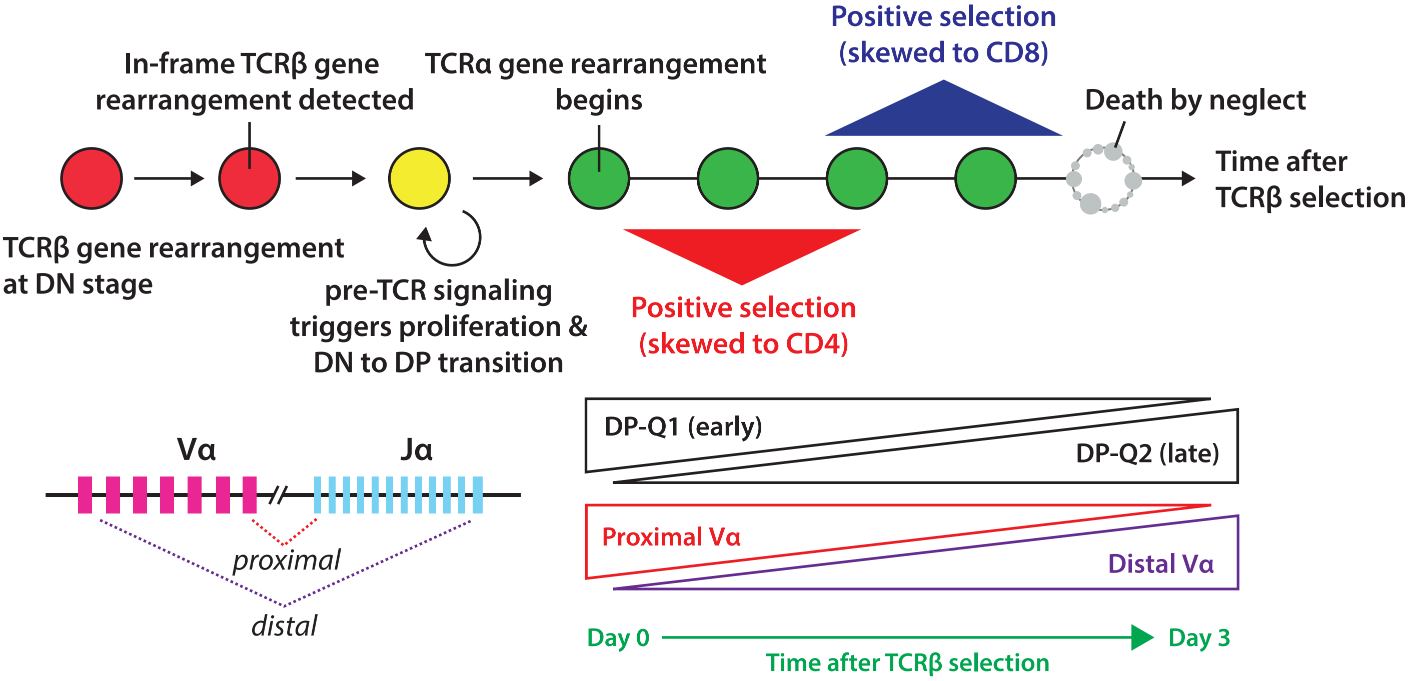
